# Sex- and strain-dependent effects of ageing on sleep and activity patterns in *Drosophila*

**DOI:** 10.1101/2023.12.30.573701

**Authors:** Nathan Woodling

## Abstract

The fruit fly *Drosophila* is a major discovery platform in the biology of ageing due to its balance of relatively short lifespan and relatively complex physiology and behaviour. Previous studies have suggested that some important phenotypes of ageing, for instance increasingly fragmented sleep, are shared from humans to *Drosophila* and can be useful measures of behavioural change with age: these phenotypes therefore hold potential as readouts of healthy ageing for genetic or pharmacological interventions aimed at the underpinning biology of ageing. However, some age-related phenotypes in *Drosophila* show differing results among studies, leading to questions regarding the source of discrepancies among experiments. In this study, I have tested females and males from three common laboratory strains of *Drosophila* to determine the extent to which sex and background strain influence age-related behavioural changes in sleep and activity patterns. Surprisingly, I find that some phenotypes – including age-related changes in total activity, total sleep, and sleep fragmentation – depend strongly on sex and strain, to the extent that some phenotypes show opposing age-related changes in different sexes or strains. Conversely, I identify other phenotypes, including age-related decreases in morning and evening anticipation, that are more uniform across sexes and strains. These results reinforce the importance of controlling for background strain in both behavioural and ageing experiments, and they imply that caution should be used when drawing conclusions from studies on a single sex or strain of *Drosophila*. At the same time, these findings also offer suggestions for behavioural measures that merit further investigation as potentially more consistent phenotypes of ageing.

## Introduction

The fruit fly *Drosophila* is one of the most widely used model organisms in ageing research: its relatively short lifespan, low maintenance cost, and extensive genetic tractability have allowed *Drosophila* research to complement work in *C. elegans* and other organisms in discovering a number of highly evolutionarily conserved genes and signalling pathways whose genetic and/or pharmacological activation or inhibition can extend lifespan (Piper & Partridge, 2018). Importantly, in many cases the lifespan-extending effects of these interventions are highly conserved from less complex species like *C. elegans* to more complex animals like mice (Campisi et al., 2019; Fontana et al., 2010; Martínez Corrales & Alic, 2020).

In addition to lifespan, healthspan (the amount of life spent in good health) is arguably an even more important outcome to consider in ageing research (Le Bourg, 2020). *Drosophila* have offered powerful tools for studying healthspan, with a prominent example in the study of sleep quality changes with age. In flies (Koh et al., 2006; Robertson & Keene, 2013), mice (Welsh et al., 1986), and humans (Bliwise, 1993; Pandi-Perumal et al., 2002; Webb, 1989), sleep is reported to become more fragmented with age, as measured by shorter individual sleep episodes or bouts. Importantly, fragmented sleep appears to be an important correlate of reduced healthspan in humans, as sleep fragmentation is predictive of both dementia and all-cause cognitive decline (Shi et al., 2018; Spira et al., 2014; Xu et al., 2020). *Drosophila* have offered additional evidence for correlations between sleep patterns and lifespan: for instance, indices of sleep fragmentation and circadian rhythmicity can significantly predict lifespan for individual flies, albeit with relatively low predictive power (Koudounas et al., 2012). *Drosophila* studies have also shown that lifespan-extending interventions can reduce age-related sleep fragmentation: for instance, deletion of genes encoding insulin-like peptides or treatment with the TOR inhibitor rapamycin can both extend lifespan (Bjedov et al., 2010; Grönke et al., 2010) and reduce age-related sleep fragmentation in flies (Metaxakis et al., 2014); similar effects of increased lifespan and reduced sleep fragmentation have been observed for inhibition of the receptor tyrosine kinase Alk in neurons (Woodling et al., 2020). Taken together, these findings suggest that age-related changes in sleep and activity patterns can be one useful behavioural measure of healthspan in *Drosophila*.

At the same time, several important caveats exist for these studies. Firstly, many studies, including those discussed above (Bjedov et al., 2010; Grönke et al., 2010; Woodling et al., 2020), have shown sex-specific or sex-biased effects of lifespan-extending interventions in *Drosophila* (Lushchak et al., 2023), suggesting that some biological mechanisms of ageing may be sex-specific. Secondly, many studies of *Drosophila* ageing are done within a single laboratory strain as a common genetic background – a practice that allows for careful control of confounding factors from genetic background, but also a practice that often limits studies to a single background strain and may thus limit the universality of findings. A careful analysis of behavioural ageing among strains and between sexes will therefore be a useful starting point from which future studies on *Drosophila* lifespan and healthspan can build. To that end, I have here investigated whether several commonly used measures of sleep and activity patterns show similar age-related trajectories in female and male flies from three different laboratory genetic background strains. My results show a surprising degree of divergence among sexes and strains for many age-related phenotypes, while highlighting some more consistent age-related behavioural changes that may guide future studies.

## Results

### Lifespan diverges among Drosophila strains and sexes

To begin investigating differences in lifespan and behaviour among strains and sexes, I chose three genetic background stocks (hereafter ‘strains’) of *Drosophila* commonly used in studies of sleep and/or ageing (see **Material and Methods**): *Canton-S* (*CS*), a wild-type (red-eyed) stock commonly used in behaviour research; *Dahomey* (*Dah*), a wild-type (red-eyed) population commonly used in ageing research; and a lab stock containing a mutant *white* allele (*w^1118^*), a stock commonly used as a background for genetic experiments involving transgenes marked by the presence of a ‘mini-white’ gene to produce coloured eyes. For each strain, I designed experiments using females and males that had been given the opportunity to mate with each other for 48 hours before being separated into single-sex conditions, to mimic the conditions commonly used in experiments designed to investigate ageing (Piper & Partridge, 2016).

I first assessed the lifespan of *CS*, *Dah*, and *w^1118^*mated female and male flies under standard physiological conditions. Within each strain, I observed that males consistently showed shorter median lifespan than females, albeit with a significant difference in the proportional lifespan difference between females and males among strains (**Figure 1**, Cox Proportional Hazards p=0.00036 for interaction between sex and strain). For instance, *CS* males showed a comparatively modest 7.4% decrease in median lifespan compared to *CS* females, whereas *Dah* and *w^1118^* males showed more substantial 30.6% and 23.1% decreases in median lifespan respectively. Interestingly, the more isogenic *CS* strain also showed more uniform ages at death with ‘steeper’ survival curves, particularly compared with the outbred *Dah* strain that has been maintained in large populations since its original collection from the wild. Taken together, these data indicate that not only absolute lifespan but also the degree to which sex modulates lifespan varies significantly among *Drosophila* background strains.

**Figure 1.**
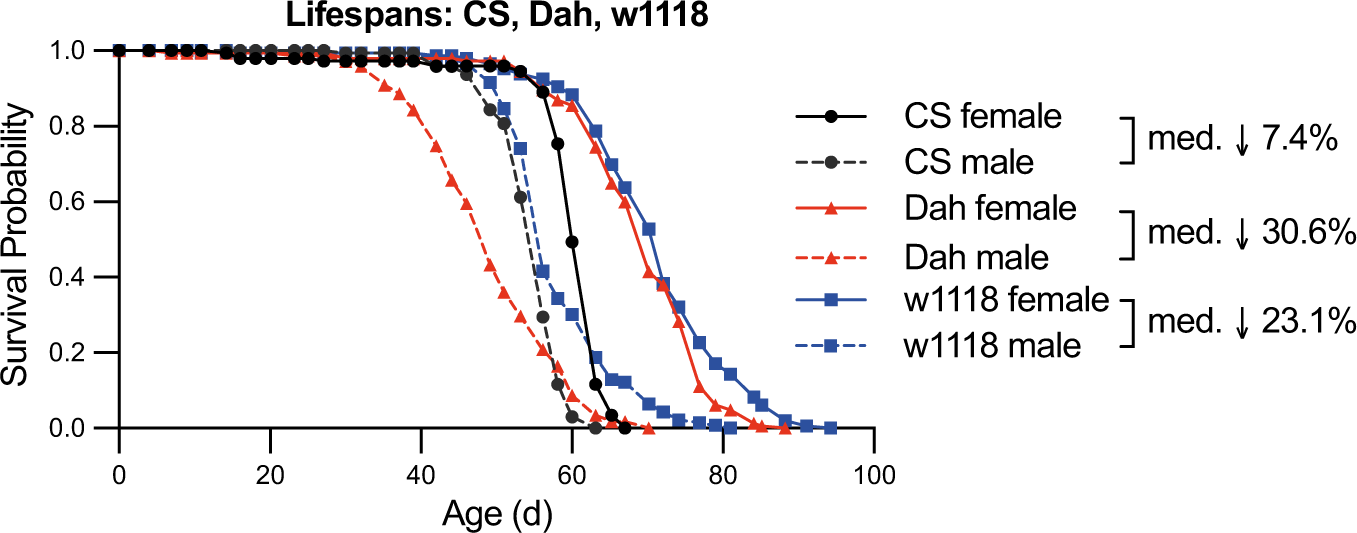
**Lifespan profiles of *CS*, *Dah*, and *w^1118^* female and male flies**. Survival curves for each sex and strain show shorter median lifespan for male flies compared with female flies within each strain, with percent decreases in median lifespan listed. Cox Proportional Hazards analysis for the interaction between sex and strain finds p=0.00036. For each pairwise comparison between sexes within a strain, log-rank tests find p<10^-20^. For each pairwise comparison between strains within a sex, log-rank tests find p<10^-6^, except for *Dah* female vs *w^1118^*female (p=0.016) and *Dah* male versus *CS* male (p=0.00049). n>125 deaths counted per condition.

### Age-related changes in activity levels depend on strain and sex

I next assessed how behavioural activity levels during the light (day) and dark (night) cycles change with age among strains and sexes. I collected mated female and male flies of each strain at 2, 4, 6, or 8 weeks of age and used the Trikinetics *Drosophila* Activity Monitor (DAM5H) system to collect activity counts per minute in 12h:12h light:dark (LD) conditions.

Plotting the mean activity counts in each 30-minute window of the 24-hour LD cycle (**Figure 2A-F**) showed strikingly divergent effects of age on behaviour among strains and sexes.

**Figure 2.**
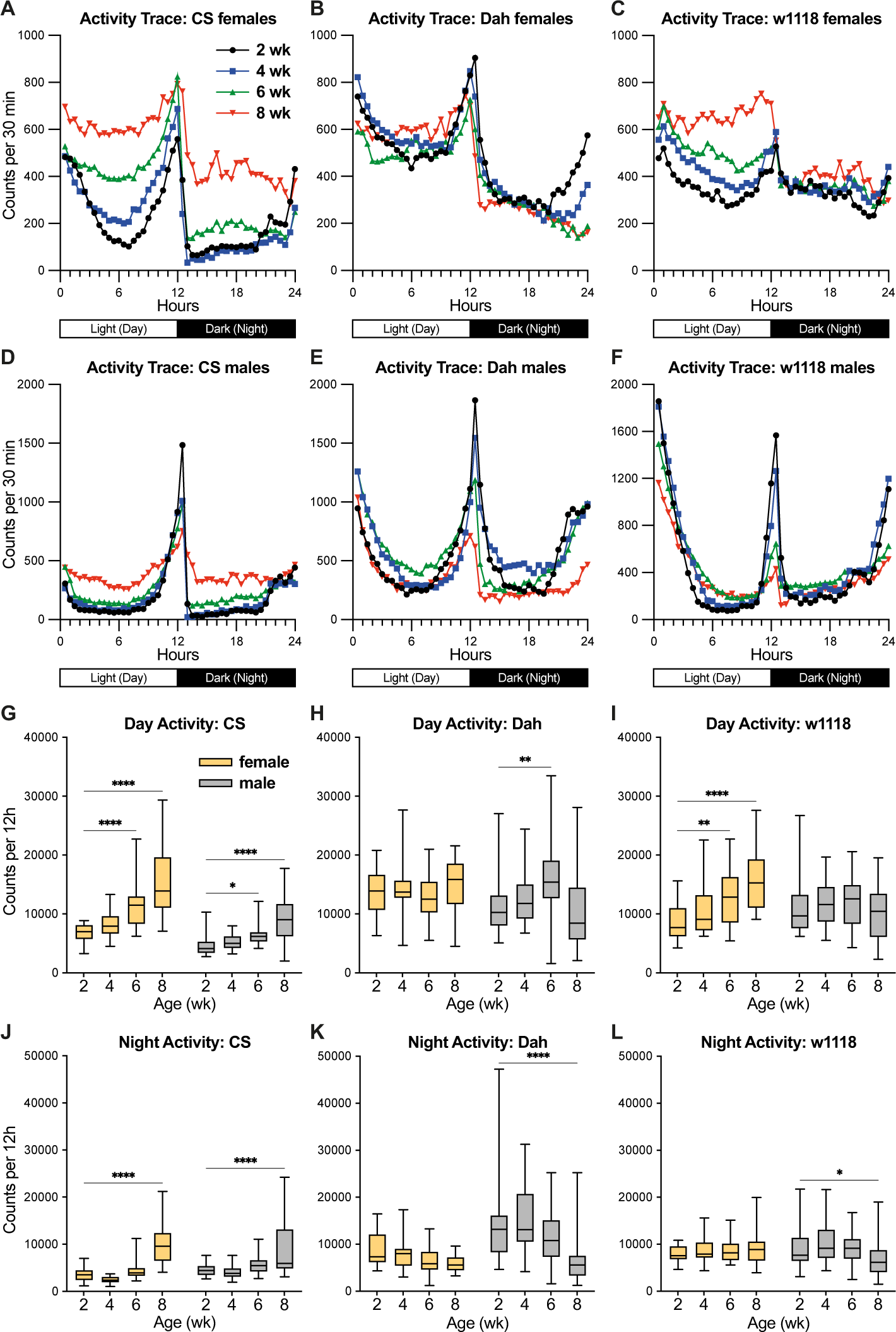
**Activity profiles of *CS*, *Dah*, and *w^1118^* female and male flies**. (**A-F**) Activity traces show the population mean of activity counts for each 30-minute bin across day and night cycles. (**G-L**) Box-and-whisker plots (minimum, 25%, median, 75%, maximum) show the summed activity counts during the day and night cycles (mean across recording days) for flies of each strain, sex, and age. n=17-32 flies per condition; * p<0.05, ** p<0.01, *** p <0.001, **** p<0.0001 by Dunnett’s multiple comparisons test versus 2-week-old group. 2-way ANOVA results are reported in **Table S1**.

Surprisingly, several populations showed significantly increased activity levels in 6-week and/or 8-week-old flies compared to 2-week-old flies (**Figure 2G-L**). Specifically, aged *CS* female and male flies showed significant increases in activity during both day and night cycles (**Figure 2G** and **2J**), whereas *w^1118^* females showed significant increases in day, but not night, activity (**Figure 2I** and **2L**). In contrast to previous studies showing moderate reductions in activity for aged CS flies (Koh et al., 2006), the only populations showing any age-related reduction in activity were *Dah* and *w^1118^*males, which showed reduced night, but not day, activity (**Figure 2K-L**). While potentially unexpected, these data indicate that age-related reduction in activity level is not a universally shared phenotype among strains and sexes.

### Age-related changes in dawn anticipation are shared among strains and between sexes

In addition to overall activity levels, day/night patterns of activity have previously been reported to change with age (Curran et al., 2019; Luo et al., 2012). *Drosophila melanogaster* individuals generally show a crepuscular pattern of activity, with higher activity levels near dusk and dawn. This pattern includes ‘anticipation’ that can be measured by an increase in activity in the hours immediately preceding the lights-on or lights-off transitions. Consistent with previous reports (Curran et al., 2019; Luo et al., 2012), I observed that young populations of flies show robust patterns of anticipation, with activity peaks preceding the lights-on morning and lights-off evening transitions (**Figure 3A**); the strength of these anticipation patterns can be quantified using published anticipation index metrics (Harrisingh et al., 2007).

**Figure 3.**
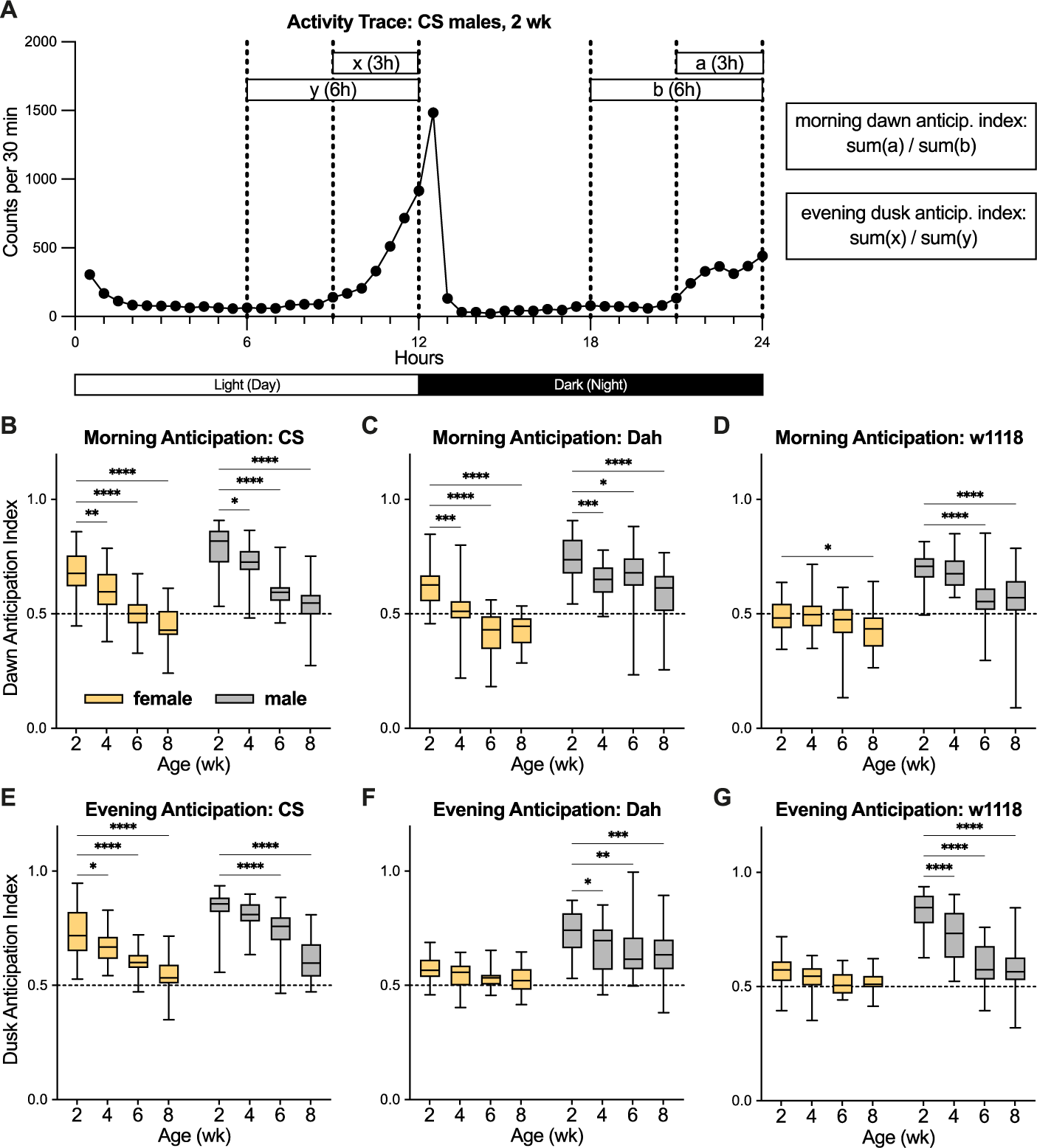
**Morning and evening anticipation indices in *CS*, *Dah*, and *w^1118^* female and male flies**. (**A**) Example activity trace (reproduced from Figure 2D) shows the method by which morning (dawn) and evening (dusk) anticipation indices are calculated. (**B-G**) Box-and-whisker plots (minimum, 25%, median, 75%, maximum) show the morning or evening anticipation indices (mean across recording days) for flies of each strain, sex, and age. n=17-32 flies per condition; * p<0.05, ** p<0.01, *** p <0.001, **** p<0.0001 by Dunnett’s multiple comparisons test versus 2-week-old group. 2-way ANOVA results are reported in **Table S1**.

When quantifying the morning and evening anticipation indices for each population of flies among my experiments, I observed that every sex and strain showed significant decreases in the morning anticipation index by 8 weeks of age, with *CS* and *Dah* females and males showing significant decreases starting even at 4 weeks of age compared to 2 weeks (**Figure 3B-D**). While I did not see as universal a pattern in age-related changes for the evening anticipation index, I observed significant reductions in this measure by 6 weeks of age for *CS* females and males, *Dah* males, and *w^1118^* males (**Figure 3E-G**). These results are particularly striking when considered in light of the strong divergence in age-related changes in overall activity levels among sexes and strains (**Figure 2**). Despite marked differences in whether populations became more or less active with age, each population consistently showed an age-related decline in their ability to anticipate the morning lights-on transition as measured by their morning anticipation index.

### Age-related changes in sleep quantity depend on strain and sex

In addition to activity patterns, sleep quantity and quality have been useful metrics of behavioural ageing in *Drosophila* (Brown et al., 2014; Koh et al., 2006; Vienne et al., 2016). To assess sleep quantity in my experimental populations, I used a widely accepted proxy measure of sleep in *Drosophila* activity assays, namely five or more consecutive minutes with zero activity counts (Huber et al., 2004; Shaw et al., 2000). When plotting the average amount of sleep in each 30-minute window of the 24-hour LD cycle (**Figure 4A-F**), I observed very striking differences among populations. For instance, males from all strains, as well as *CS* females, showed sleep patterns consistent with a large body of published work: these flies tended to show high wakefulness in the morning, move to a period of increased sleep in the middle of the day, then show high wakefulness again in the evening before a more consolidated period of sleep at night (**Figure 4A** and **4D-F**). In contrast, *Dah* and *w^1118^* females showed very little sleep compared to other populations, with *w^1118^* females showing little clear pattern of sleep between day and night cycles (**Figure 4B-C**). When quantifying age-related changes in sleep amount (**Figure 4G-L**), I also observed striking differences among strains and sexes. Potentially consistent with the age-related hyperactivity I observed in *CS* females and males (**Figure 2G** and **2J**), I observed significant age-related decreases in sleep amount in *CS* females and males by 6 weeks of age (**Figure 4G** and **4J**). At the same time, this correlation between age-related changes in activity levels and age-related sleep amount was not shared among all populations: for instance, aged *w^1118^* males showed no clear change in daytime activity levels with increasing age (**Figure 2I**), but this same population showed strongly significant age-related decreases in daytime sleep with increasing age (**Figure 4I**). These findings indicate that, while sleep quantity offers a distinct behavioural measure from activity levels, neither measure shows consistent changes with age across the strains and sexes in these experiments.

**Figure 4.**
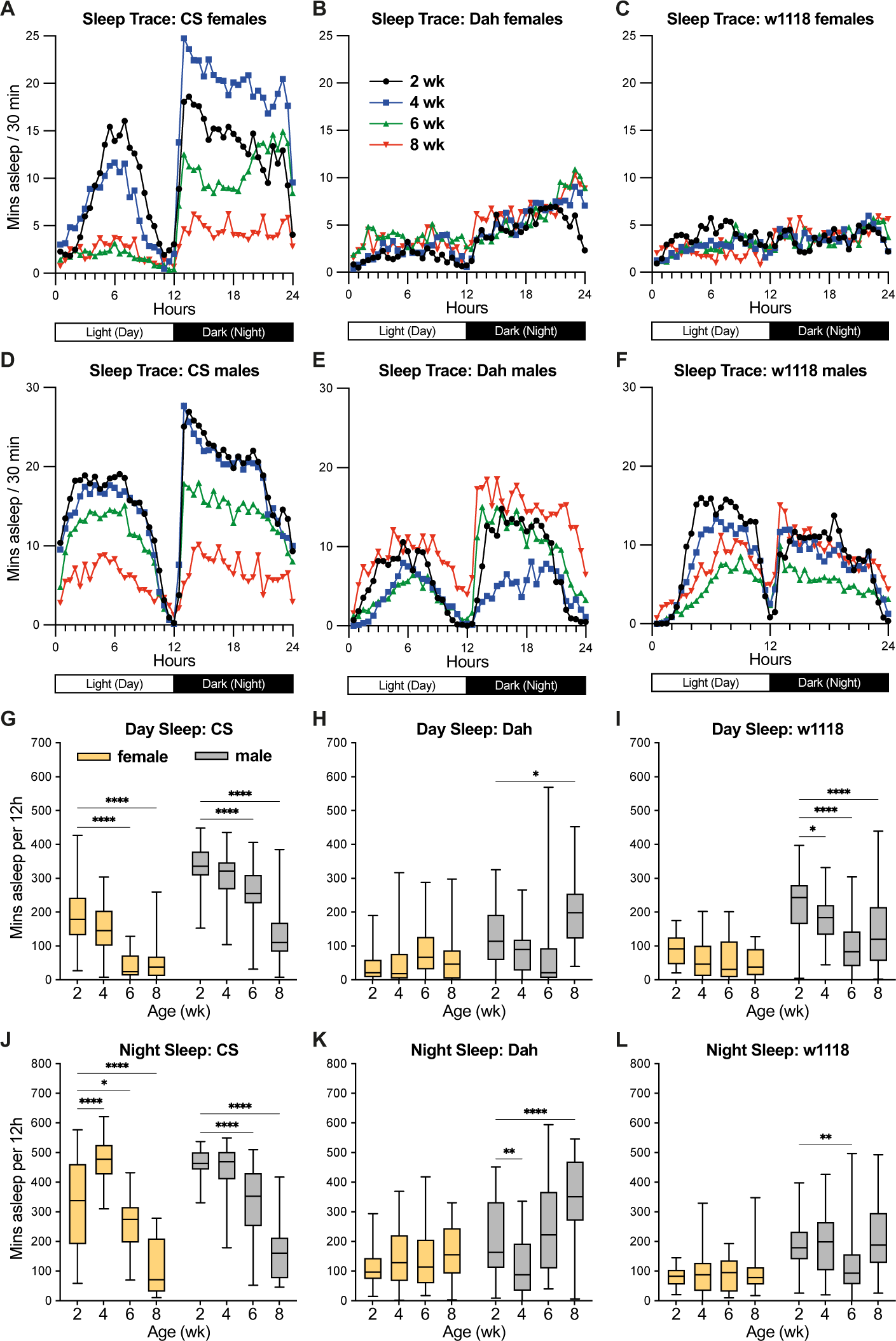
**Sleep profiles of *CS*, *Dah*, and *w^1118^* female and male flies**. (**A-F**) Sleep traces show the population mean of sleep minutes (defined as 5 or more consecutive minutes of 0 activity counts) for each 30-minute bin across day and night cycles. (**G-L**) Box-and-whisker plots (minimum, 25%, median, 75%, maximum) show the summed sleep minutes during the day and night cycles (mean across recording days) for flies of each strain, sex, and age. n=17-32 flies per condition; * p<0.05, ** p<0.01, **** p<0.0001 by Dunnett’s multiple comparisons test versus 2- week-old group. 2-way ANOVA results are reported in **Table S1**.

### Age-related changes in sleep quality depend on strain and sex

Sleep quality – namely the way sleep is consolidated into longer bouts (individual sleep episodes) or fragmented into shorter bouts – has been one of the most widely studied measures of age-related behavioural change in *Drosophila*, with studies generally reporting that older flies have higher numbers of sleep bouts with shorter sleep bout duration (indicating more fragmented sleep), particularly during the night (Brown et al., 2014; Koh et al., 2006; Vienne et al., 2016). To assess sleep quality among the experimental populations here, I first assessed the mean number of day and night sleep bouts among ages, strains, and sexes (**Figure 5**). Here, I observed no consistent pattern among strains and sexes: for instance, *CS* females and males displayed decreasing numbers of sleep bouts with increasing age (**Figure 5A** and **5D**), whereas *Dah* males displayed increased numbers of sleep bouts at the oldest ages (**Figure 5B** and **5E**).

**Figure 5.**
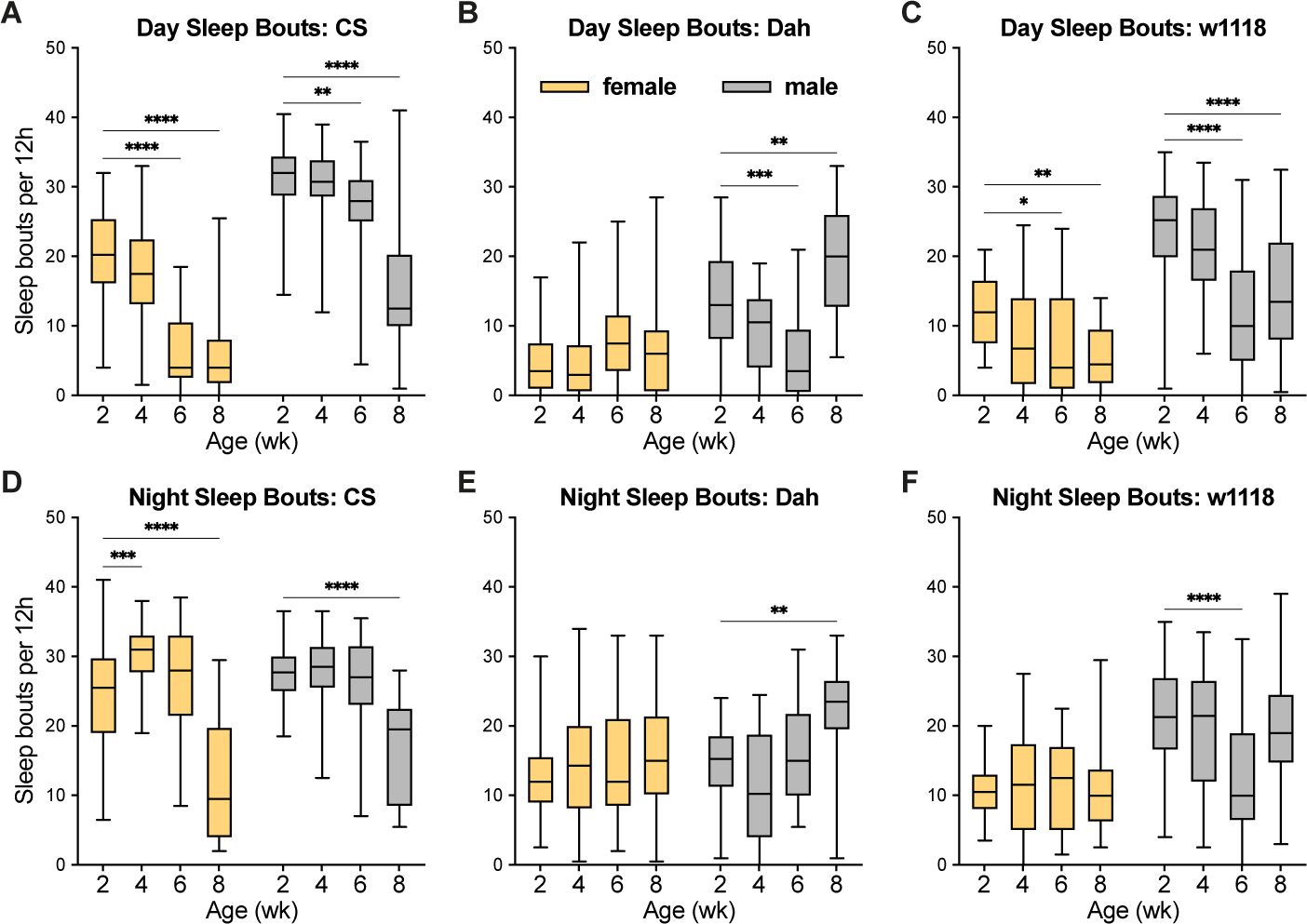
**Sleep bout numbers in *CS*, *Dah*, and *w^1118^* female and male flies**. (**A-F**) Box-and- whisker plots (minimum, 25%, median, 75%, maximum) show the number of independent sleep bouts (defined as 5 or more consecutive minutes of 0 activity counts) during the day and night cycles (mean across recording days) for flies of each strain, sex, and age. n=17-32 flies per condition; * p<0.05, ** p<0.01, *** p <0.001, **** p<0.0001 by Dunnett’s multiple comparisons test versus 2-week-old group. 2-way ANOVA results are reported in **Table S1**.

To consider another measure of sleep quality, I next assessed the mean sleep bout length among ages, strains, and sexes (**Figure 6**). Here, I observed some patterns consistent with previously published studies: for instance, 6-week and 8-week-old *CS* females and males all showed shorter sleep bout length during both the day and night as compared to 2-week-old flies (**Figure 6A** and **6D**). However, I did not observe a similar pattern in either *Dah* or *w1118* flies, suggesting a high level of divergence in how sleep quality changes with age among *Drosophila* background strains. Taken together with my data on sleep quantity (**Figure 4**), these analyses do not suggest that sleep patterns change with age in a consistent way among strains and sexes; instead, they suggest that the measures of sleep and activity most relevant for ageing may depend heavily on factors such as background strain and sex.

**Figure 6.**
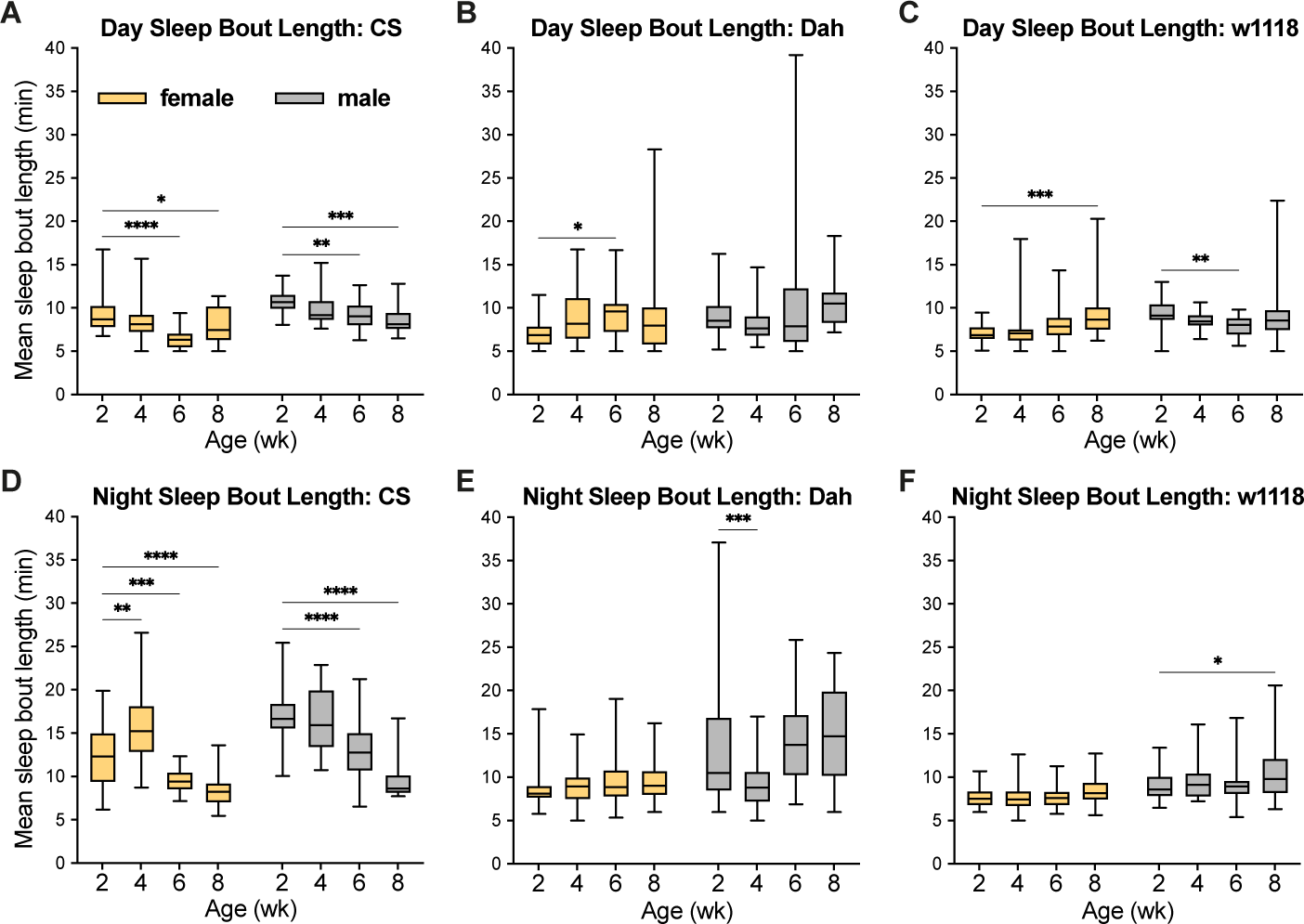
**Sleep bout length in *CS*, *Dah*, and *w^1118^* female and male flies**. (**A-F**) Box-and- whisker plots (minimum, 25%, median, 75%, maximum) show the length of sleep bouts (defined as 5 or more consecutive minutes of 0 activity counts) during the day and night cycles (mean across recording days) for flies of each strain, sex, and age. n=17-32 flies per condition; * p<0.05, ** p<0.01, *** p <0.001, **** p<0.0001 by Dunnett’s multiple comparisons test versus 2-week-old group. 2-way ANOVA results are reported in **Table S1**.

## Discussion

Increasingly, research on ageing using model organisms has considered both lifespan (as measured by survival) and healthspan (as measured by changes in behaviour and other phenotypes with age) (Le Bourg, 2020). *Drosophila* has been a productive model system for investigating sex differences in both lifespan and healthspan, as well as the sex-specific plasticity of these phenotypes with longevity-promoting interventions (Regan et al., 2016, 2022). With respect to lifespan, my current results are consistent with many of these past studies: in each strain investigated, I observed significantly greater longevity in female compared to male flies (**Figure 1**). However, the extent to which females outlived males differed greatly among strains, suggesting that studies investigating sex differences in ageing may come to slightly different conclusions depending on the background strain used. Importantly, my lifespan results differ from one of the foundational studies on age-related sleep and activity changes in *Drosophila*, where the authors observed longer lifespan in male than in female *CS* flies (Koh et al., 2006).

This suggests that additional factors, for example diet and specific rearing conditions, likely contribute to sex differences in lifespan.

With respect to healthspan, my data only partially overlap with previous studies, and they highlight some important differences among strains and sexes. Firstly, my data on age-dependent sleep fragmentation (as measured by decreased sleep bout length) parallel only a subset of results from previous studies. For instance, one of the first studies to thoroughly investigate age-related sleep change in flies found partially overlapping results with mine: they found that *CS* virgin female and male flies show a transient increase in sleep bout length in early life, followed by a decline in sleep bout length in late life (Koh et al., 2006). These data closely parallel my own results for mated female and male *CS* flies, but they are markedly different from my results for *w^1118^* and *Dah* flies of both sexes (**Figure 6**). In addition, I observe strong age-related increases in activity counts for *CS* flies (**Figure 2**) which were not observed in the previous study (Koh et al., 2006). Other more recent studies have examined both *CS* and *w^1118^*flies and found that both strains exhibited age-dependent declines in sleep bout length (Brown et al., 2014; Vienne et al., 2016), as has another study examining virgin female flies of the *w^Dah^* strain (in which the *w^1118^* allele has been backcrossed into an otherwise *Dah* genetic background) (Metaxakis et al., 2014). However, yet other studies have failed to observe age-dependent shortening of sleep bout length in male flies derived from a *CS* background (Bushey et al., 2010). Taken together, these results highlight sex- and strain-dependent discrepancies among previous literature and my own results on age-dependent increases in sleep fragmentation.

At the same time, my data highlight other behavioural phenotypes, such as anticipation indices, that more closely align with previous findings and show robust age-dependent changes across sexes and strains. Previous work has shown that *w^1118^-iso^31^* female and male flies show age-dependent reductions in both morning and evening anticipation indices (Curran et al., 2019; Luo et al., 2012), mirroring my own results for these measures in almost all strains and sexes examined here (**Figure 3**). Notably, my data suggest that morning anticipation is more robustly age-dependent than evening anticipation, again mirroring previous findings (Curran et al., 2019). As noted in this previous work, morning anticipation behaviour is controlled by a very specific subset of neurons – the ventral lateral neurons – that express the neuropeptide Pdf (Grima et al., 2004; Stoleru et al., 2004). Moreover, Pdf levels decrease with age in *CS* males (Umezaki et al., 2012), and RNAi-based knockdown of *Pdf* in ventral lateral neurons reduces morning anticipation in young male flies (Shafer & Taghert, 2009). Taken together, these findings present an attractive hypothesis in which age-dependent loss of Pdf in a specific class of neurons directly leads to age-dependent decline in morning anticipation. When coupled with my current findings of declining morning anticipation in females and males from three different strains, these data suggest that reduced Pdf expression and a resulting decline in morning anticipation have the potential to be robust phenotypes of ageing that might occur broadly across sexes and strains.

What factors might contribute to the discrepancies between the current results and previous studies for some but not all phenotypes? One major difference may lie in the nature of the activity monitors used: many previous studies have used the Trikinetics DAM2 system in which a single infrared beam passes through each single-fly tube, whereas the data in the current study are derived from the newer generation Trikinetics DAM5H system in which 15 infrared beams pass through each tube. In theory, the DAM5H system should allow for finer-scale movement detection even when flies move within a small region of the tube – meaning that these data will likely report higher activity and lower sleep levels than previous studies. However, phenotypes in which data are normalised to activity level (such as the anticipation indices that employ a ratio rather than absolute numbers) may be more reproducible between DAM systems – perhaps explaining why these measures show more similar age-dependent trajectories between previous studies and the current work. As one potential downside of the DAM5H system, some users have reported spurious activity counts from oversensitive detectors, even in empty tubes; however, in experiments where I included a set of empty tubes in parallel with the fly studies shown here, I was not able to detect any spurious counts in my experimental setup (n=120 beams, 8 tubes; 0 counts for all beams across >90 hours of recording).

Additional factors may also contribute to seemingly discordant findings between this work and previous studies. *Drosophila* diet, including the specific type of each ingredient, can have major effects on age-related phenotypes (Bass et al., 2007); even the specific lot number of yeast from the same company can alter *Drosophila* phenotypes, as can the type of preservative used (Sannino & Dobson, 2023). The *Drosophila* diet used in the current study is very similar to some previous studies on age-related sleep changes (Metaxakis et al., 2014) but distinct from others (Brown et al., 2014; Curran et al., 2019; Koh et al., 2006; Luo et al., 2012; Vienne et al., 2016); diet may therefore be an additional factor explaining differences among studies. Sex and mating status can also have profound effects on age-related phenotypes in *Drosophila*: for instance, females display higher levels of gut hyperplasia and permeability with age (Hudry et al., 2016; Regan et al., 2016) in a manner dependent on both mating status (Ahmed et al., 2020) and background strain (Regan et al., 2016). My current results may therefore compare most easily with other studies using mated female and male flies rather than those that have used virgin flies. Importantly, my results also find significant age-by-sex interactions for all measures of behaviour in at least one of the strains tested here (**Table S1**); it is therefore highly likely that behavioural studies will uncover different or even opposing effects of anti-ageing interventions in females versus males. Finally, while this work has used three different strains in an attempt to find robust phenotypes across *Drosophila*, this is still a relatively limited number of strains and will not reflect the full range of age-related behavioural changes among the large number of common *Drosophila* lab strains.

Given the strong divergence of age-related behavioural change among strains and sexes, what conclusions can be drawn from existing and future studies using these measures as correlates of healthspan? Firstly, and as a note of moderation, the current data should not necessarily call into question previous studies using *Drosophila* to examine age-related sleep or activity changes. Well-designed studies have offered important molecular mechanistic insight into genetic or pharmacological interventions that extend lifespan and delay behavioural features of ageing like sleep fragmentation within a strain and/or sex (Metaxakis et al., 2014). Future investigation into whether these mechanisms extend to other strains and sexes will therefore be important follow-up research to understand how universal these mechanisms are. Secondly, the current results suggest that some measures of behaviour – for example morning or evening anticipation – could potentially offer more reproducible measures of age-related behavioural change among strains and between sexes. Finally, these results reinforce the importance of backcrossing and other strategies to control for background strain in studies examining mutant or transgenic animals, as the background strain of each mutant or transgenic line may differ greatly in behaviour and therefore lead to spurious results. Control of genetic background has long been recognised as an important benchmark for research on ageing (Partridge & Gems, 2007) and other physiological quantitative traits (Evangelou et al., 2019); the current data therefore underscore the importance of proper controls in genetic studies. Importantly, background strain effects are likely to affect studies in mammals as well, as age-related behavioural changes can also be strain-dependent in mice (Hasan et al., 2012). Ultimately, these results suggest a note of caution should be used when drawing conclusions from even carefully controlled behavioural studies on single sexes or background strains in any model organism.

## Materials and Methods

### Fly stocks and husbandry

*Drosophila* stocks were maintained and experiments conducted at 25°C on a 12h:12h light:dark cycle on food containing 10% (w/v) brewer’s yeast (MP Bio, lot number U1122284494-1), 5% (w/v) sucrose, 1.5% (w/v) agar, 0.3% (w/v) Nipagin, and 0.3% (v/v) propionic acid.

The *Canton-S* (*CS*) wild-type stock was originally isogenised in 1943 (Stern & Schaeffer, 1943) and has been widely used in neurogenetics research since being chosen by Seymour Benzer for its homogeneous phototactic behaviour (Benzer, 1967); the *CS* stock used here was a kind gift from the lab of Colin McClure (Queen’s University Belfast). The wild-caught *Dahomey (Dah)* wild-type stock was collected in 1970 in Dahomey (now Benin) and has since been maintained in large populations with overlapping generations; the *Dah* stock used here was obtained from the lab of Linda Partridge (University College London). The *w^1118^* stock used here was also obtained from the Partridge lab, where it has been used as a background for neurodegenerative disease model research (Sofola et al., 2010) and maintained in smaller population sizes. All stocks were maintained under identical conditions (multiple large population bottles) for at least three generations before being used in the present studies.

### Survival analysis

Lifespan assays were carried out as described in detail in (Piper & Partridge, 2016). From the eggs collected from each set of parental flies, the progeny that emerged as adults within a 24- hour window were collected and allowed to mate for 48 hours, after which they were separated into single-sex vials containing standard food at a density of 15 individuals per vial. Flies were transferred to fresh vials three times per week, with deaths and censors scored during each transfer. Microsoft Excel (template available at http://piperlab.org/resources/) was used to calculate survival proportions after each transfer.

### Drosophila Activity Monitor experiments

Flies were generated and maintained as for survival analysis above, with successive groups of mated female and male flies generated at 2-week intervals. When the oldest cohort of flies for a strain reached 8 weeks of age, all four cohorts (2-week, 4-week, 6-week, and 8-week) were briefly anaesthetised and transferred to single-fly tubes (Trikinetics, 65mm glass tubes) containing a small amount of standard fly food at one end and a small cotton plug at the other end. These tubes were then inserted into DAM5H Drosophila Activity Monitors (DAMs, Trikinetics) with the food adjacent to beam number ‘1’. DAMs were then placed in an incubator at 25°C on a 12h:12h light:dark cycle for at least 72 hours (sufficient time to gather data before any fertilised eggs produced larvae that could confound activity counts). Data were collected using the DAMSystem3 software (Trikinetics) with activity counts collected once each minute. All flies were allowed at least 24 hours to acclimatise to the DAMs; data were then extracted for 48 hours starting from the next lights-on event. During these 48 hours of data recording, the exact ages of each cohort were 13-14 days, 27-28 days, 41-42 days, and 55-56 days.

Data on activity counts were analysed using R (scripts available at Mendley Data, doi: 10.17632/8633hm46p5.1) in RStudio Version 2023.03.1+446. These scripts were designed to reproduce previous Excel-based analysis pipelines described in (Chen et al., 2019). Briefly, the first script translated Trikinetics monitor files containing per-beam counts to files containing total activity counts summed over all beams. The second script calculated per-fly measures of activity counts (total, day, and night); anticipation indices (morning and evening); sleep amounts (total, day, and night); sleep bout number (total, day, and night); and sleep bout length (total, day, and night). For all experiments, sleep was defined as 5 or more consecutive minutes of inactivity, as widely defined in the literature (Huber et al., 2004; Shaw et al., 2000). Morning and evening anticipation indices were calculated as shown in **Figure 3A**, as previously defined (Harrisingh et al., 2007). This script also excluded any flies that died during the recording; for these experiments, deaths were defined as flies with fewer than 3 activity counts during the final 180 minutes of analysed recording. The third script compiled data from across different monitor files to create a single table for each measure across experimental conditions. Analysed data were then exported to GraphPad Prism 10.0 for graphical display and statistical analysis.

### Statistical analysis

The graphical displays (generated in GraphPad Prism 10.0) and statistical tests used for each readout are described in the associated figure legend. Log-rank tests were performed in Microsoft Excel (template available at http://piperlab.org/resources/); Cox Proportional Hazards tests were performed in R using the ‘survival’ package; and 2-way ANOVA with Dunnett’s multiple comparisons tests were performed in GraphPad Prism 10.0. For all statistical tests, p<0.05 was considered significant.

### Data availability

All raw data and metadata underlying each figure can be freely obtained at Mendeley Data, doi: 10.17632/8633hm46p5.1.

## Acknowledgements

I am grateful for the diligent and professional work from the staff of the University of Glasgow Academic Service Unit in their preparation of the *Drosophila* media needed to run these experiments. I also thank the labs of Adam Dobson and Alberto Sanz Montero for equipment sharing and scientific advice throughout this project. I gratefully acknowledge the Royal Society (Research Grant RGS\R1\231248) for funding this work.

## Author Contributions

Conceptualisation, Methodology, Investigation, Formal analysis, Writing, Visualisation, Project administration, and Funding acquisition: N.W.

## Open Access Statement

For the purpose of open access in line with University of Glasgow policy, the author has applied a Creative Commons Attribution (CC BY) licence to any Author Accepted Manuscript version arising from this submission.

## Competing Interests

I declare that I have no competing interests.

---

**Table S1.**
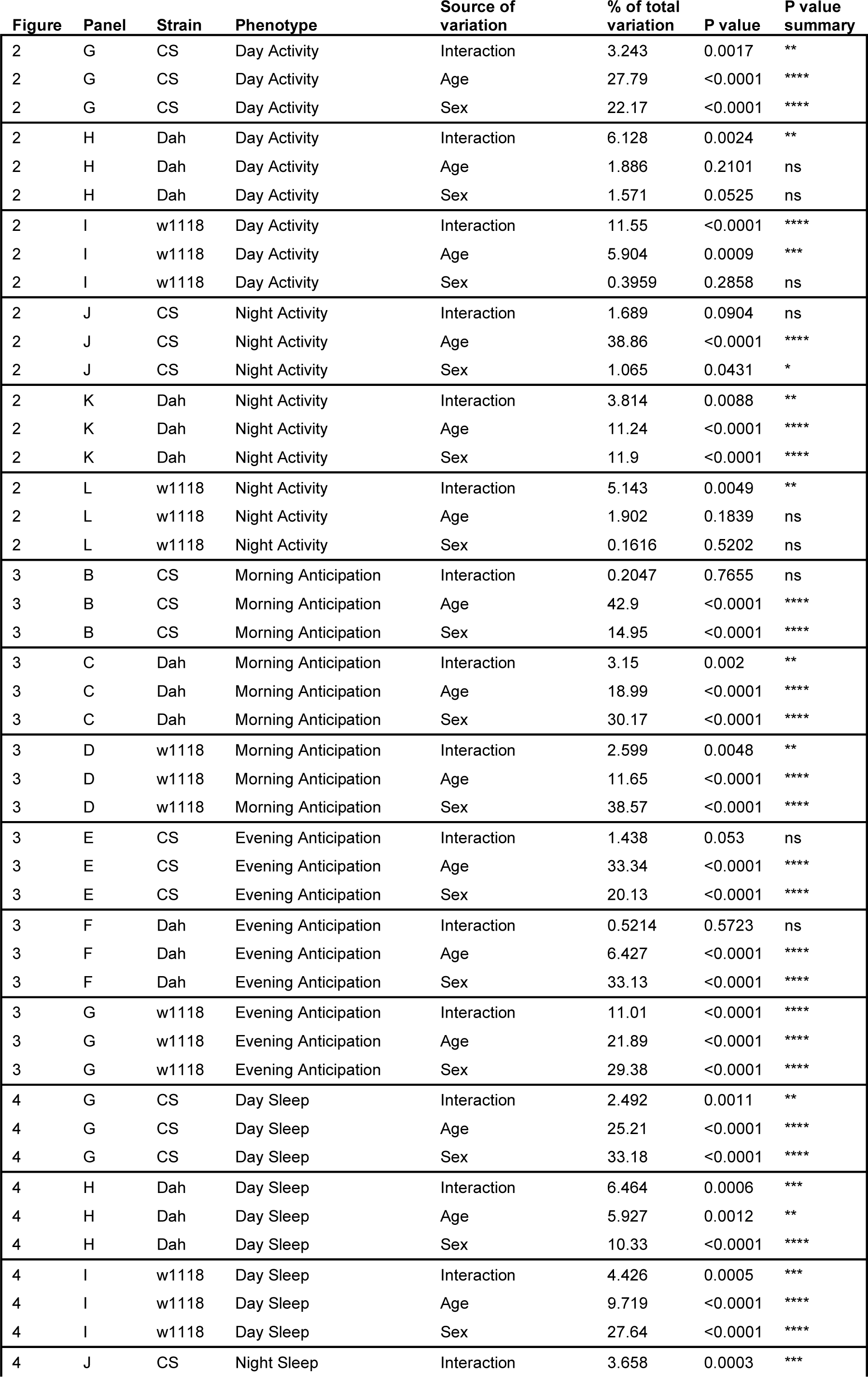
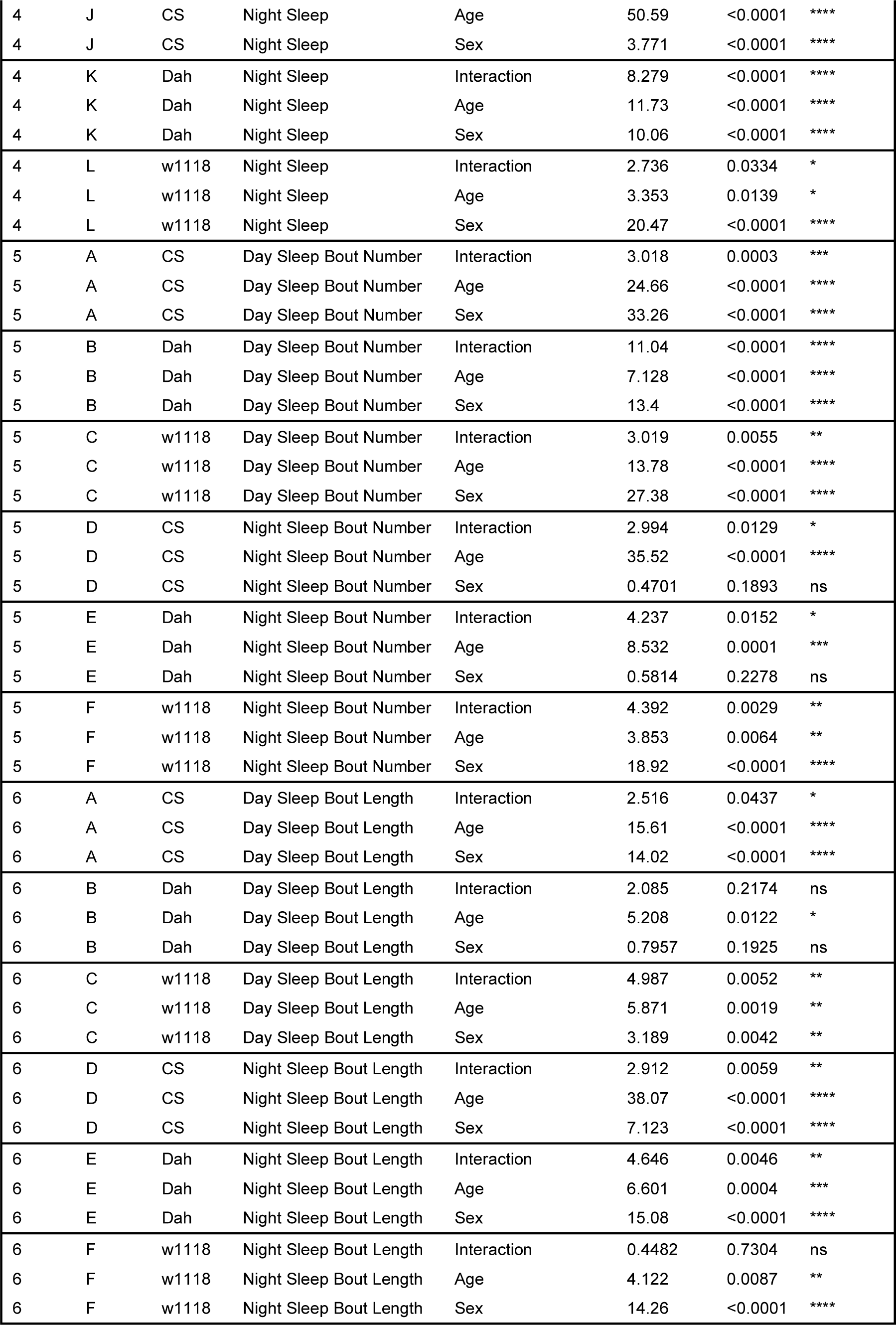
2-way ANOVA results for all figures.

## Notes

### Competing Interest Statement

The authors have declared no competing interest.

## References

1. Ahmed, S. M. H., Maldera, J. A., Krunic, D., Paiva-Silva, G. O., Pénalva, C., Teleman, A. A., & Edgar, B. A. (2020). Fitness trade-offs incurred by ovary-to-gut steroid signalling in Drosophila. Nature 2020 *584*:7821, *584*(7821), 415–419. 10.1038/s41586-020-2462-y

2. Bass, T. M., Grandison, R. C., Wong, R., Martinez, P., Partridge, L., & Piper, M. D. W. (2007). Optimization of dietary restriction protocols in Drosophila. Journals of Gerontology - Series A Biological Sciences and Medical Sciences, 62(10), 1071–1081. 10.1093/gerona/62.10.1071

3. Benzer, S. (1967). Behavioral mutants of Drosophila isolated by countercurrent distribution. Proceedings of the National Academy of Sciences, 58(3), 1112–1119. 10.1073/pnas.58.3.1112

4. Bjedov, I., Toivonen, J. M., Kerr, F., Slack, C., Jacobson, J., Foley, A., & Partridge, L. (2010). Mechanisms of Life Span Extension by Rapamycin in the Fruit Fly Drosophila melanogaster. Cell Metabolism, 11(1), 35–46. 10.1016/j.cmet.2009.11.010

5. Bliwise, D. L. (1993). Sleep in Normal Aging and Dementia. Sleep, 16(1), 40–81. 10.1093/SLEEP/16.1.40

6. Brown, M. K., Chan, M. T., Zimmerman, J. E., Pack, A. I., Jackson, N. E., & Naidoo, N. (2014). Aging induced endoplasmic reticulum stress alters sleep and sleep homeostasis. Neurobiology of Aging, 35(6), 1431–1441. 10.1016/J.NEUROBIOLAGING.2013.12.005

7. Bushey, D., Hughes, K. A., Tononi, G., & Cirelli, C. (2010). Sleep, aging, and lifespan in Drosophila. BMC Neuroscience, 11(1), 1–18. 10.1186/1471-2202-11-56/TABLES/2

8. Campisi, J., Kapahi, P., Lithgow, G. J., Melov, S., Newman, J. C., & Verdin, E. (2019). From discoveries in ageing research to therapeutics for healthy ageing. Nature, 571(7764), 183–192. 10.1038/s41586-019-1365-2

9. Chen, K. F., Lowe, S., Lamaze, A., Krätschmer, P., & Jepson, J. (2019). Neurocalcin regulates nighttime sleep and arousal in Drosophila. ELife, 8, 1717. 10.7554/eLife.38114

10. Curran, J. A., Buhl, E., Tsaneva-Atanasova, K., & Hodge, J. J. L. (2019). Age-dependent changes in clock neuron structural plasticity and excitability are associated with a decrease in circadian output behavior and sleep. Neurobiology of Aging, 77, 158. 10.1016/J.NEUROBIOLAGING.2019.01.025

11. Evangelou, A., Ignatiou, A., Antoniou, C., Kalanidou, S., Chatzimatthaiou, S., Shianiou, G., Ellina, S., Athanasiou, R., Panagi, M., Apidianakis, Y., & Pitsouli, C. (2019). Unpredictable Effects of the Genetic Background of Transgenic Lines in Physiological Quantitative Traits. G3: Genes|Genomes|Genetics, 9(11), 3877. 10.1534/G3.119.400715

12. Fontana, L., Partridge, L., & Longo, V. D. (2010). Extending healthy life span-from yeast to humans. Science, 328(5976), 321–326. 10.1126/science.1172539

13. Grima, B., Chélot, E., Xia, R., & Rouyer, F. (2004). Morning and evening peaks of activity rely on different clock neurons of the Drosophila brain. Nature 2004 *431*:7010, *431*(7010), 869–873. 10.1038/nature02935

14. Grönke, S., Clarke, D. F., Broughton, S., Andrews, T. D., & Partridge, L. (2010). Molecular evolution and functional characterization of Drosophila insulin-like peptides. PLoS Genetics, 6(2), e1000857. 10.1371/journal.pgen.1000857

15. Harrisingh, M. C., Wu, Y., Lnenicka, G. A., & Nitabach, M. N. (2007). Intracellular Ca2+ Regulates Free-Running Circadian Clock Oscillation In Vivo. Journal of Neuroscience, 27(46), 12489–12499. 10.1523/JNEUROSCI.3680-07.2007

16. Hasan, S., Dauvilliers, Y., Mongrain, V., Franken, P., & Tafti, M. (2012). Age-related changes in sleep in inbred mice are genotype dependent. Neurobiology of Aging, 33(1), 195.e13- 195.e26. 10.1016/J.NEUROBIOLAGING.2010.05.010

17. Huber, R., Hill, S. L., Holladay, C., Biesiadecki, M., Tononi, G., & Cirelli, C. (2004). Sleep Homeostasis in Drosophila Melanogaster. Sleep, 27(4), 628–639. 10.1093/SLEEP/27.4.628

18. Hudry, B., Khadayate, S., & Miguel-Aliaga, I. (2016). The sexual identity of adult intestinal stem cells controls organ size and plasticity. Nature, 530(7590), 344–348. 10.1038/nature16953

19. Koh, K., Evans, J. M., Hendricks, J. C., & Sehgal, A. (2006). A Drosophila model for age-associated changes in sleep:wake cycles. Proceedings of the National Academy of Sciences, 103(37), 13843–13847. 10.1073/pnas.0605903103

20. Koudounas, S., Green, E. W., & Clancy, D. (2012). Reliability and variability of sleep and activity as biomarkers of ageing in Drosophila. Biogerontology, 13(5), 489–499. 10.1007/S10522-012-9393-4/FIGURES/6

21. Le Bourg, E. (2020). Lifespan Versus Healthspan (pp. 439–452). Springer, Cham. 10.1007/978-3-030-52663-4_25

22. Luo, W., Chen, W. F., Yue, Z., Chen, D., Sowcik, M., Sehgal, A., & Zheng, X. (2012). Old flies have a robust central oscillator but weaker behavioral rhythms that can be improved by genetic and environmental manipulations. Aging Cell, 11(3), 428–438. 10.1111/J.1474-9726.2012.00800.X

23. Lushchak, O., Strilbytska, O., & Storey, K. B. (2023). Gender-specific effects of pro-longevity interventions in Drosophila. Mechanisms of Ageing and Development, 209, 111754. 10.1016/J.MAD.2022.111754

24. Martínez Corrales, G., & Alic, N. (2020). Evolutionary Conservation of Transcription Factors Affecting Longevity. Trends in Genetics, 36(5), 373–382. 10.1016/j.tig.2020.02.003

25. Metaxakis, A., Tain, L. S., Grönke, S., Hendrich, O., Hinze, Y., Birras, U., & Partridge, L. (2014). Lowered Insulin Signalling Ameliorates Age-Related Sleep Fragmentation in Drosophila. PLoS Biology, 12(4), e1001824. 10.1371/journal.pbio.1001824

26. Pandi-Perumal, S. R., Seils, L. K., Kayumov, L., Ralph, M. R., Lowe, A., Moller, H., & Swaab, D. F. (2002). Senescence, sleep, and circadian rhythms. Ageing Research Reviews, 1(3), 559–604. 10.1016/S1568-1637(02)00014-4

27. Partridge, L., & Gems, D. (2007). Benchmarks for ageing studies. Nature, 450(7167), 165–167. 10.1038/450165a

28. Piper, M. D. W., & Partridge, L. (2016). Protocols to Study Aging in Drosophila. In Methods in Molecular Biology (Vol. 1478, Issue Chapter 18, pp. 291–302). Springer New York. 10.1007/978-1-4939-6371-3_18

29. Piper, M. D. W., & Partridge, L. (2018). Drosophila as a model for ageing. Biochimica et Biophysica Acta (BBA) - Molecular Basis of Disease, 1864(9), 2707–2717. 10.1016/J.BBADIS.2017.09.016

30. Regan, J. C., Khericha, M., Dobson, A. J., Bolukbasi, E., Rattanavirotkul, N., & Partridge, L. (2016). Sex difference in pathology of the ageing gut mediates the greater response of female lifespan to dietary restriction. ELife, 5, e10956. 10.7554/eLife.10956

31. Regan, J. C., Lu, Y.-X., Ureña, E., Meilenbrock, R. L., Catterson, J. H., Kißler, D., Fröhlich, J., Funk, E., & Partridge, L. (2022). Sexual identity of enterocytes regulates autophagy to determine intestinal health, lifespan and responses to rapamycin. Nature Aging, 2(12), 1145– 1158. 10.1038/s43587-022-00308-7

32. Robertson, M., & Keene, A. C. (2013). Molecular Mechanisms of Age-Related Sleep Loss in the Fruit Fly - A Mini-Review. Gerontology, 59(4), 334–339. 10.1159/000348576

33. Sannino, D. R., & Dobson, A. J. (2023). Acetobacter pomorum in the Drosophila gut microbiota buffers against host metabolic impacts of dietary preservative formula and batch variation in dietary yeast. Applied and Environmental Microbiology, 89(10), e0016523. 10.1128/aem.00165-23

34. Shafer, O. T., & Taghert, P. H. (2009). RNA-Interference Knockdown of Drosophila Pigment Dispersing Factor in Neuronal Subsets: The Anatomical Basis of a Neuropeptide’s Circadian Functions. PLOS ONE, 4(12), e8298. 10.1371/JOURNAL.PONE.0008298

35. Shaw, P. J., Cirelli, C., Greenspan, R. J., & Tononi, G. (2000). Correlates of sleep and waking in Drosophila melanogaster. *Science (New York*, N.Y.), 287(5459), 1834–1837. 10.1126/science.287.5459.1834

36. Shi, L., Chen, S. J., Ma, M. Y., Bao, Y. P., Han, Y., Wang, Y. M., Shi, J., Vitiello, M. V., & Lu, L. (2018). Sleep disturbances increase the risk of dementia: A systematic review and meta-analysis. Sleep Medicine Reviews, 40, 4–16. 10.1016/J.SMRV.2017.06.010

37. Sofola, O., Kerr, F., Rogers, I., Killick, R., Augustin, H., Gandy, C., Allen, M. J., Hardy, J., Lovestone, S., & Partridge, L. (2010). Inhibition of GSK-3 ameliorates Aβ pathology in an adult-onset Drosophila model of Alzheimer’s disease. PLoS Genetics, 6(9), e1001087. 10.1371/journal.pgen.1001087

38. Spira, A. P., Chen-Edinboro, L. P., Wu, M. N., & Yaffe, K. (2014). Impact of sleep on the risk of cognitive decline and dementia. Current Opinion in Psychiatry, 27(6), 478–483. 10.1097/YCO.0000000000000106

39. Stern, C., & Schaeffer, E. W. (1943). On Wild-Type Iso-Alleles in Drosophila Melanogaster. Proceedings of the National Academy of Sciences, 29(11), 361–367. 10.1073/PNAS.29.11.361

40. Stoleru, D., Peng, Y., Agosto, J., & Rosbash, M. (2004). Coupled oscillators control morning and evening locomotor behaviour of Drosophila. Nature 2004 *431*:7010, *431*(7010), 862–868. 10.1038/nature02926

41. Umezaki, Y., Yoshii, T., Kawaguchi, T., Helfrich-Förster, C., & Tomioka, K. (2012). Pigment-dispersing factor is involved in age-dependent rhythm changes in Drosophila melanogaster. Journal of Biological Rhythms, 27(6), 423–432. 10.1177/0748730412462206

42. Vienne, J., Spann, R., Guo, F., & Rosbash, M. (2016). Age-Related Reduction of Recovery Sleep and Arousal Threshold in Drosophila. Sleep, 39(8), 1613–1624. 10.5665/SLEEP.6032

43. Webb, W. B. (1989). Age-related Changes in Sleep. Clinics in Geriatric Medicine, 5(2), 275– 287. 10.1016/S0749-0690(18)30678-5

44. Welsh, D. K., Richardson, G. S., & Dement, W. C. (1986). Effect of Age on the Circadian Pattern of Sleep and Wakefulness in the Mouse. Journal of Gerontology, 41(5), 579–586. 10.1093/GERONJ/41.5.579

45. Woodling, N. S., Aleyakpo, B., Dyson, M. C., Minkley, L. J., Rajasingam, A., Dobson, A. J., Leung, K. H. C., Pomposova, S., Fuentealba, M., Alic, N., & Partridge, L. (2020). The neuronal receptor tyrosine kinase Alk is a target for longevity. Aging Cell, 19(5), e13137. 10.1111/acel.13137

46. Xu, W., Tan, C. C., Zou, J. J., Cao, X. P., & Tan, L. (2020). Original research: Sleep problems and risk of all-cause cognitive decline or dementia: an updated systematic review and meta-analysis. *Journal of Neurology*, Neurosurgery, and Psychiatry, 91(3), 236. 10.1136/JNNP-2019-321896

